# SpaIM: Single-cell Spatial Transcriptomics Imputation via Style Transfer

**DOI:** 10.1101/2025.01.24.634756

**Authors:** Bo Li, Ziyang Tang, Aishwarya Budhkar, Xiang Liu, Tonglin Zhang, Baijian Yang, Jing Su, Qianqian Song

## Abstract

Spatial transcriptomics (ST) technologies have revolutionized our understanding of cellular ecosystems. However, these technologies face challenges such as sparse gene signals and limited gene detection capacities, which hinder their ability to fully capture comprehensive spatial gene expression profiles. To address these limitations, we propose leveraging single-cell RNA sequencing (scRNA-seq), which provides comprehensive gene expression data but lacks spatial context, to enrich ST profiles. Herein, we introduce SpaIM, an innovative style transfer learning model that utilizes scRNA-seq information to predict unmeasured gene expressions in ST data, thereby improving gene coverage and expressions. SpaIM segregates scRNA-seq and ST data into data-agnostic contents and data-specific styles, with the contents capture the commonalities between the two data types, while the styles highlight their unique differences. By integrating the strengths of scRNA-seq and ST, SpaIM overcomes data sparsity and limited gene coverage issues, making significant advancements over 12 existing methods. This improvement is demonstrated across 53 diverse ST datasets, spanning sequencing- and imaging-based spatial technologies in various tissue types. Additionally, SpaIM enhances downstream analyses, including the detection of ligand-receptor interactions, spatial domain characterization, and identification of differentially expressed genes. Released as open-source software, SpaIM increases accessibility for spatial transcriptomics analysis. In summary, SpaIM represents a pioneering approach to enrich spatial transcriptomics using scRNA-seq data, enabling precise gene expression imputation and advancing the field of spatial transcriptomics research.

## INTRODUCTION

Recent advances in spatial transcriptomics (ST) technologies have emerged to provide deep insights into spatial cellular ecosystems^1-3^. Sequencing-based ST technologies^4-7^, such as 10x Genomics Visium^8^ and Slide-seq^9, 10^, utilize spatially indexed barcodes to conduct RNA sequencing on tissue spots. Meanwhile, imaging-based ST platforms like NanoString’s CosMx™ SMI^11^ and Vizgen’s MERSCOPE^12^ employ in situ hybridization and fluorescence microscopy to provide spatial transcriptomics data at the single-cell level. Despite these advancements, the gene expression profiles from these ST technologies exhibit data sparsity and limited gene coverage. For instance, NanoString’s CosMx™ SMI^11^ only assays thousands of genes, and the actual number of mRNA molecules detected per cell remains low, resulting in poor gene expression measurement due to limitations in molecular imaging and hybridization efficiency. This inherent technological constraint limits both the comprehensiveness of gene coverage and the density of count data, posing significant challenges. Addressing these limitations through computational methods is crucial to fully capturing and interpreting spatial transcriptomics profiles.

Before the advent of spatial transcriptomics, single-cell RNA sequencing (scRNA-seq) technologies have gained attention for their ability to elucidate cellular heterogeneity^13-16^ and trace cell lineages^17-19^. Despite such insights, scRNA-seq lacks spatial information, making it challenging to determine the structural organization of cells within complex tissues. Nonetheless, as a complement to ST data, scRNA-seq has become invaluable for enhancing the quality of spatial transcriptomics, facilitating precise analyses of the transcriptome with spatial resolution in individual tissue sections. To improve spatial transcriptomics profiles, researchers have been actively developing methods^20-27^ to seamlessly integrate scRNA-seq with ST data. Notable methods include Tangram^20^, gimVI^22^, and spaGE^21^. Specifically, Tangram^20^ uses regularizers to filter an optimal subset of scRNA-seq profiles mapping with the spatial data. gimVI^22^ adopts a deep generative model to integrate scRNA-seq and ST data for the imputation of missing genes. SpaGE^21^ utilizes Principal Component Analysis to identify principal vectors and align cells from scRNA-seq and ST by k-nearest-neighbor. novoSpaRc^25^ leverages the continuity of gene expression among neighboring cells for predictions. Recent methods such as stDiff^28^ and SpatialScope^29^ use deep generative models to impute spatial gene expressions. TISSUE^30^ and SPRITE^31^ use uncertainty-aware and meta-approach to achieve spatial gene expression predictions. Other methods like Seurat^23^, SpaOTsc^24^, LIGER^26^, and stPlus^27^ utilize different computational strategies to achieve local alignments between scRNA-seq and ST data, enabling the prediction of unmeasured gene expressions in ST data. However, these existing methods have inherent limitations as they primarily rely on local alignment to predict unmeasured gene expressions, which cannot fully unleash the potential of scRNA-seq and ST data for gene expression prediction.

In this study, we introduce SpaIM, i.e. Spatial transcriptomics IMputation, a novel style transfer learning^32^ framework that leverages scRNA-seq data to impute unmeasured or missing gene expressions in ST data. SpaIM consists of an ST autoencoder and an ST generator, which work together to decouple scRNA-seq data and ST data into data-agnostic contents and data-specific styles. The data-agnostic contents capture the shared information between scRNA-seq and ST data, while the data-specific styles reflect the intrinsic differences between scRNA-seq and ST data. After training with a specifically designed loss function, the ST generator can independently predict unmeasured gene expressions in ST data using only scRNA-seq data, ensuring accurate imputation. SpaIM is available as open-source software on GitHub (https://github.com/QSong-github/SpaIM), with detailed tutorials demonstrating its capabilities in enhancing the utility of spatial transcriptomics profiles.

## RESULTS

### Overview of the SpaIM model

To accurately impute gene expressions including unmeasured ones in spatial transcriptomics (ST) data, we introduce SpaIM, a novel style transfer learning model leveraging scRNA-seq (SC) data. As illustrated in **Fig.1a**, SpaIM is a multilayer Recursive Style Transfer (ReST) model with layer-wise content- and style-based feature extraction and fusion. Specifically, SpaIM comprises an ST autoencoder (**Fig.1a**) and an ST generator (**Fig.1a**) that are constructed with ReST. For a single gene, we consider the gene expression pattern across the *K* single cell clusters as its content, and the unique gene expression pattern across all cells in ST data, which differs from SC data, as its style. The style represents the intrinsic differences in gene expressions between the ST and the SC platforms. The style-transfer learning of SpaIM involves two simultaneous tasks: the ST autoencoder uses the SC data as the reference to disentangle the ST gene expression patterns into content and style, and the ST generator extracts the content from the SC data and transfers the learned style from ST autoencoder to infer ST gene expressions. The ST autoencoder and the ST generator share the same decoder and are co-trained using a joint loss function based on the common genes between ST and SC data. This allows the ST generator to capture the gene expression patterns in the ST data as well as the relation between the ST and SC data. After training, the ST generator is used as a stand-alone model to infer the expression patterns of unmeasured genes in the ST data, using only the SC data as input. In this way, the well trained SpaIM model enables accurate predictions of unmeasured gene expressions, through leveraging the comprehensive gene expression profiles of scRNA-seq and the optimized ST generator.

**Fig. 1.**
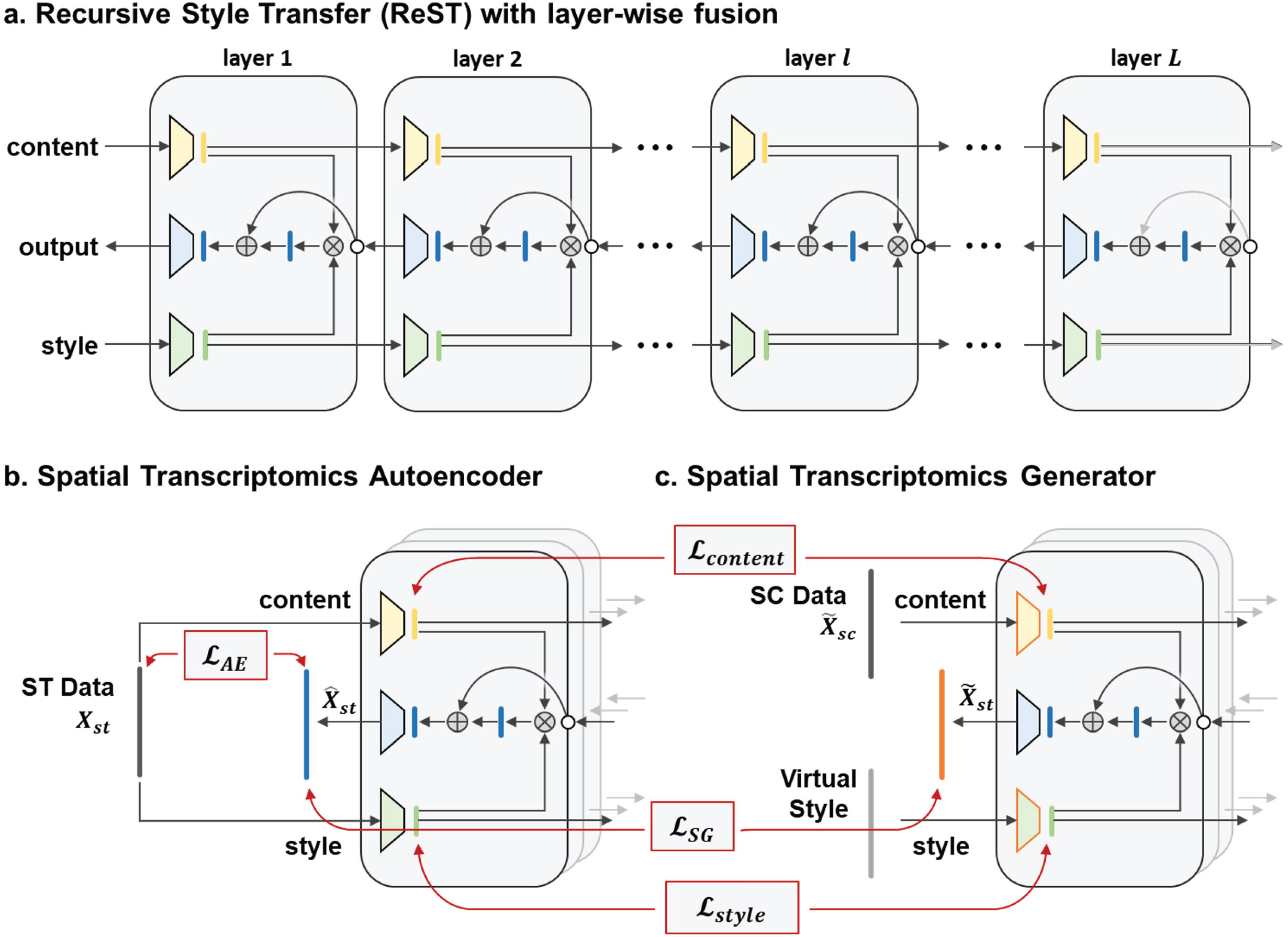
Overview of the SpaIM model. SpaIM comprises the comprises an ST autoencoder and an ST generator. Both ST autoencoder and ST generator are built on the multilayer Recursive Style Transfer (ReST) layers.

### SpaIM accurately imputes spatial gene expression in human breast cancer tissue slice

To evaluate the spatial gene imputation capabilities of our model, we conduct the performance comparisons using ST (10x Visium, ‘CID44971’) and scRNA-seq data (10x Chromium, GSE176078) from breast cancer tissues. Data source and details are listed in **Supplementary Table 1**. To comprehensively evaluate the performance of SpaIM, we compare it with 12 existing methods including Tangram^20^, SPRITE^31^, stDiff^28^, SpatialScope^29^, TISSUE^30^, gimVI^22^, SpaGE^21^, Seurat^23^, SpaOTsc^24^, novoSpaRc^25^, LIGER^26^, and stPlus^27^. Evaluation metrics (see **Materials and Methods**) include the Pearson correlation coefficient (PCC), structural similarity index measure (SSIM), Jaccard similarity (JS), root mean square error (RMSE), and accuracy score (ACC). Higher values of PCC, SSIM, and ACC, as well as lower values of JS and RMSE represent better performance.

The comparison results are shown in **Fig.2a**, which highlights the superior performance of SpaIM over other methods across all metrics. Specifically, **Fig.2a** reveals that SpaIM achieves the best values across all four metrics (PCC: 0.70±0.02), outperforming the second-best model, Tangram (PCC: 0.62±0.02). Of note, the SSIM value of SpaIM (SSIM: 0.60±0.02) shows 10% higher than that of Tangram (SSIM: 0.52±0.02, **Fig.2a**), demonstrating that the imputed gene expressions of SpaIM are much closer to the ground truth. Furthermore, **Supplementary Fig.1a** illustrates the other metrics, with SpaIM achieving a significantly better performance (JS: 0.11±0.01, RMSE: 0.81±0.01) than other methods. This underscores the exceptional accuracy of SpaIM in spatial gene expression imputation.

**Fig. 2.**
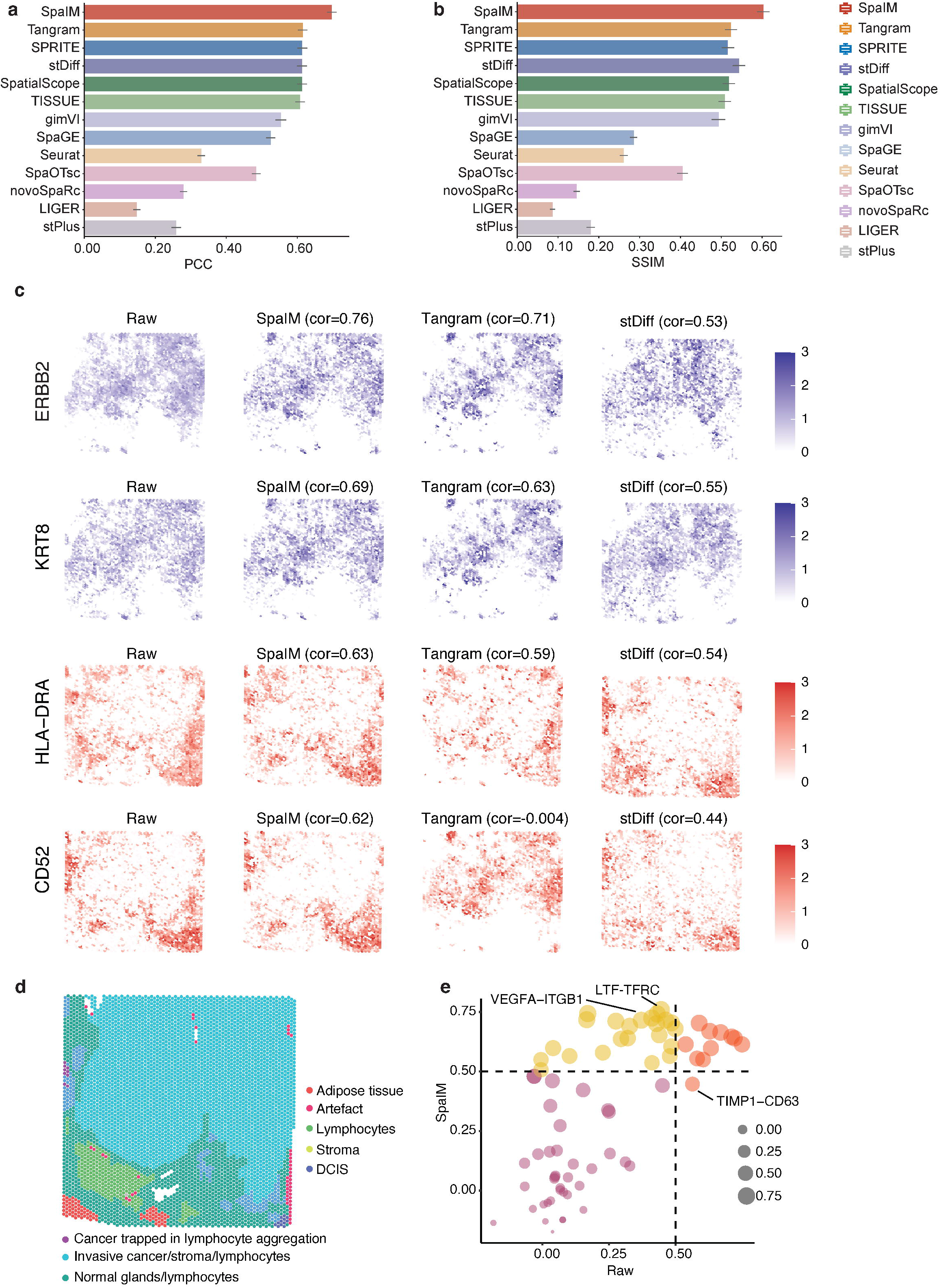
Performance evaluation in the breast cancer dataset. **a** Comparison results between SpaIM and existing methods using Pearson correlation coefficient (PCC). **b** Comparison results between SpaIM and existing methods using structural similarity index measure (SSIM). **c** Spatial visualization of the ground truth and the predicted gene expressions. **d** Spatial visualization of spatial domains. **e** Scatter plot of associated ligand □ receptor pairs in the raw data and the SpaIM-imputed data.

In addition to the comprehensive evaluations of SpaIM, we further examine the gene expression predictions generated by different methods. For this breast cancer ST data, **Fig.2a** presents the predicted gene expressions of maker genes for the invasive cancer region (*ERBB2, KRT8*, **Fig.2a**), with PCC values indicating their correlations with ground truth. SpaIM consistently outperforms other methods, surpassing both Tangram and stDiff. Moreover, SpaIM accurately predicts marker genes in the normal gland region, such as *HLA-DRA* and *CD52* (**Fig.2a**), achieving PCC of 0.63 and 0.62, respectively. In contrast, Tangram produces misleading predictions for CD52, with a notably low correlation. Next, to validate the utility of SpaIM-imputed data for downstream analysis, we curate a combined list of candidate ligand □ receptor (L-R) pairs^33^. Among the collected L-R pairs, SpaIM-imputed data identified 33 strongly associated pairs, compared to only 11 pairs detected in the raw data, with 10 pairs overlapping (**Fig.2a**), suggesting the capability of SpaIM in revealing biological insights. Top associated pairs such as VEGFA-ITGB1^34^ and LTF-TFRC^35, 36^, well known for their roles in tumor activities, are exclusively detected in the SpaIM-imputed data. Collectively, these results underscore the reliability and superiority of SpaIM in gene expression imputation.

### SpaIM enhances detection of differentially expressed genes

As an emerging imaging-based ST technology, NanoString CosMx™ enables the detection of up to a thousand genes per slide^37^ at subcellular resolution, emphasizing the necessity of using SpaIM to expand gene coverage. Here we collect CosMx ST datasets from lung cancer tissues (**Supplementary Table 1**), with up to 70k to 130k cells per slide, to evaluate the performance of SpaIM.

Taking the Lung9-rep1 dataset as an example, we assess the gene expression imputation efficacy of different methods. **Fig.3a** clearly shows that SpaIM, with median SSIM and JS values of 0.21 and 0.43, outperforms other methods such as Tangram (SSIM: 0.15, JS: 0.47), stDiff (SSIM: 0.12, JS: 0.50), SpatialScope (SSIM: 0.11, JS: 0.50), gimVI (SSIM: 0.19, JS: 0.66), SpaGE (SSIM: 0.10, JS: 0.62), and Seurat (SSIM: 0.17, JS: 0.82). Moreover, SpaIM consistently achieves superior performance across additional metrics including PCC and RMSE, than the other methods such as gimVI and Tangram (**Supplementary Fig.1a**). Notably, in terms of accuracy score (ACC), SpaIM shows significantly higher ACC (ACC: 0.96±0.05), compared to other methods including Tangram (ACC: 0.91±0.11), stDiff (ACC: 0.59±0.09), SpatialScope (ACC: 0.55±0.17), gimVI (ACC: 0.82±0.27), SpaGE (ACC: 0.64±0.14), and Seurat (ACC: 0.36±0.14).

**Fig. 3.**
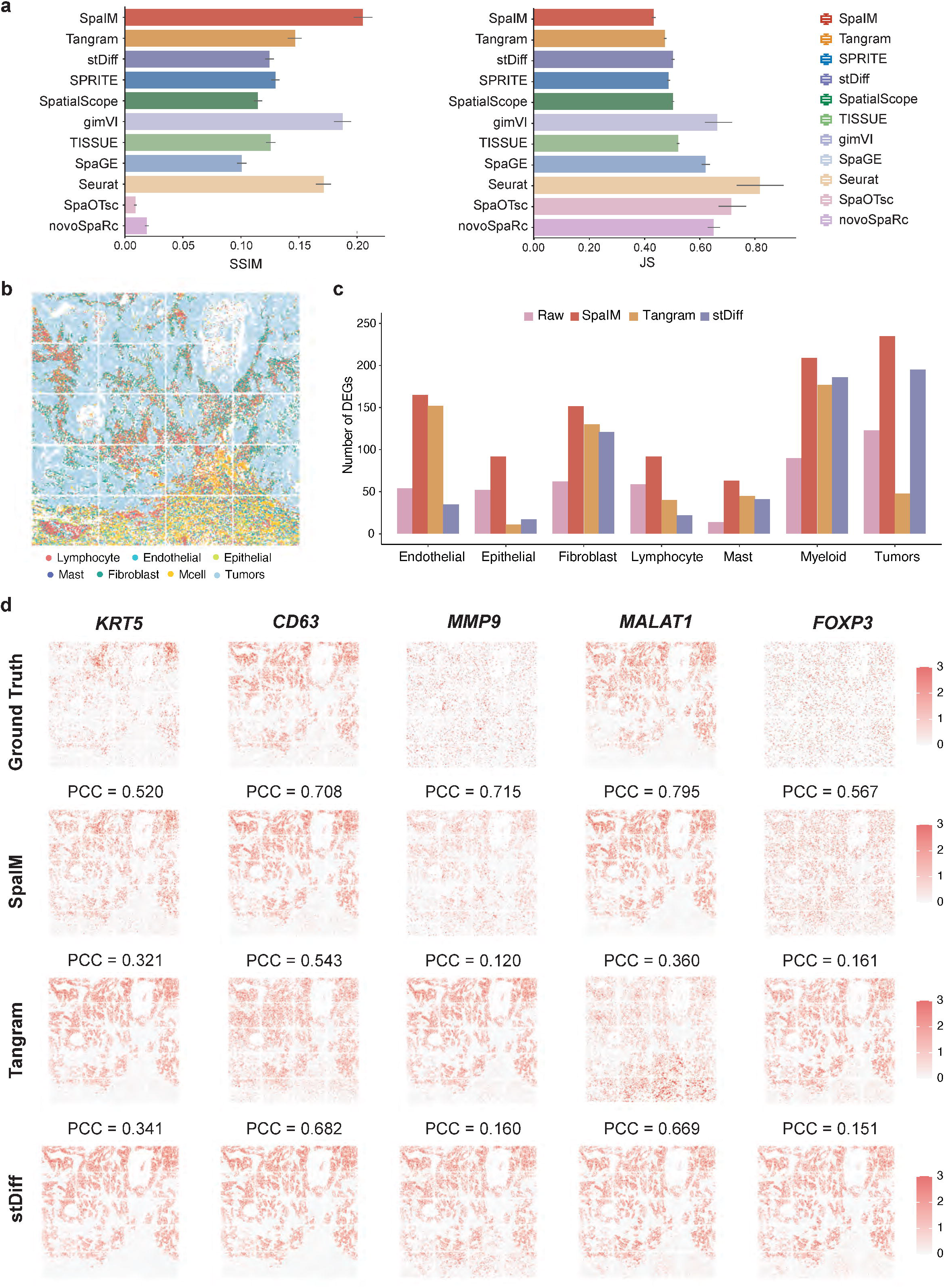
Benchmarking performance on the NanoString CosMx SMI dataset. **a** Benchmarking results on the Lung9-rep1 dataset, using evaluation metrics including structural similarity index measure (SSIM) and Jaccard similarity (JS). **b** Spatial visualization of cell types in the Lung9-rep1 dataset. **c** Comparisons of the number of differentially expressed genes in each cell type. **d** Spatial visualization of ground truth and the predicted expressions of tumor-related genes.

With different cell types in the spatial microenvironment (**Fig.3a**), we then identify the differentially expressed genes (DEGs) for each cell type using raw data and imputed data from the top-performing methods, i.e. SpaIM, Tangram, and stDiff (**Fig.3a**). As expected, compared to the 59 lymphocyte-specific DEGs detected in the raw data with an adjusted P-value of 0.05 and a log2 fold change of 1, the number of significant genes increase to 92 when using SpaIM imputations. In contrast, Tangram and stDiff imputations identify 40 and 22 DEGs, respectively, at the same thresholds. Moreover, to visually illustrate the predicted spatial gene patterns by different methods, we specifically select biologically significant DEGs (*KRT5, CD63, MMP9, MALAT1, FOXP3*) as examples to evaluate whether SpaIM can accurately recover their profiles if they are masked (i.e., considered as unmeasured genes). As is known, those genes play pivotal roles in cancer, participating in various signaling pathways that regulate tumor biological behaviors including epithelial-mesenchymal transition^38^, immune evasion^39^, tumor progression^40^, and metastasis^41, 42^. The raw and predicted gene expressions are visualized with PCC values labeled (**Fig.3a**). For intuitive comparison, we include the ground truth gene expressions in the top row. Remarkably, the spatial gene patterns predicted by SpaIM exhibit higher similarity to the ground truth, as evidenced by a superior PCC and a closely matched threshold range, underscoring SpaIM’s robust capacity for imputing biologically significant genes.

### SpaIM facilities spatial domain detection and recovers unmeasured genes

In addition, we evaluate the imputation performance on another single-cell spatial data (Lung5-rep3). Specifically, **Fig.4a** shows that SpaIM achieves a higher median SSIM of 0.15 in the Lung5-rep3 dataset, outperforming other methods including Tangram (median SSIM: 0.14), stDiff (median SSIM: 0.13), and SPRITE (median SSIM: 0.12). SpaIM also exhibits superior performance (JS: 0.36), lower than Tangram (JS: 0.49) by approximately 13% and stDiff (JS: 52) by about 16%. Other metrics including PCC, RMSE, and accuracy (**Supplementary Fig.1a**) further demonstrate the outperformance of SpaIM, establishing it as the most effective method for imputing gene expressions.

**Fig. 4.**
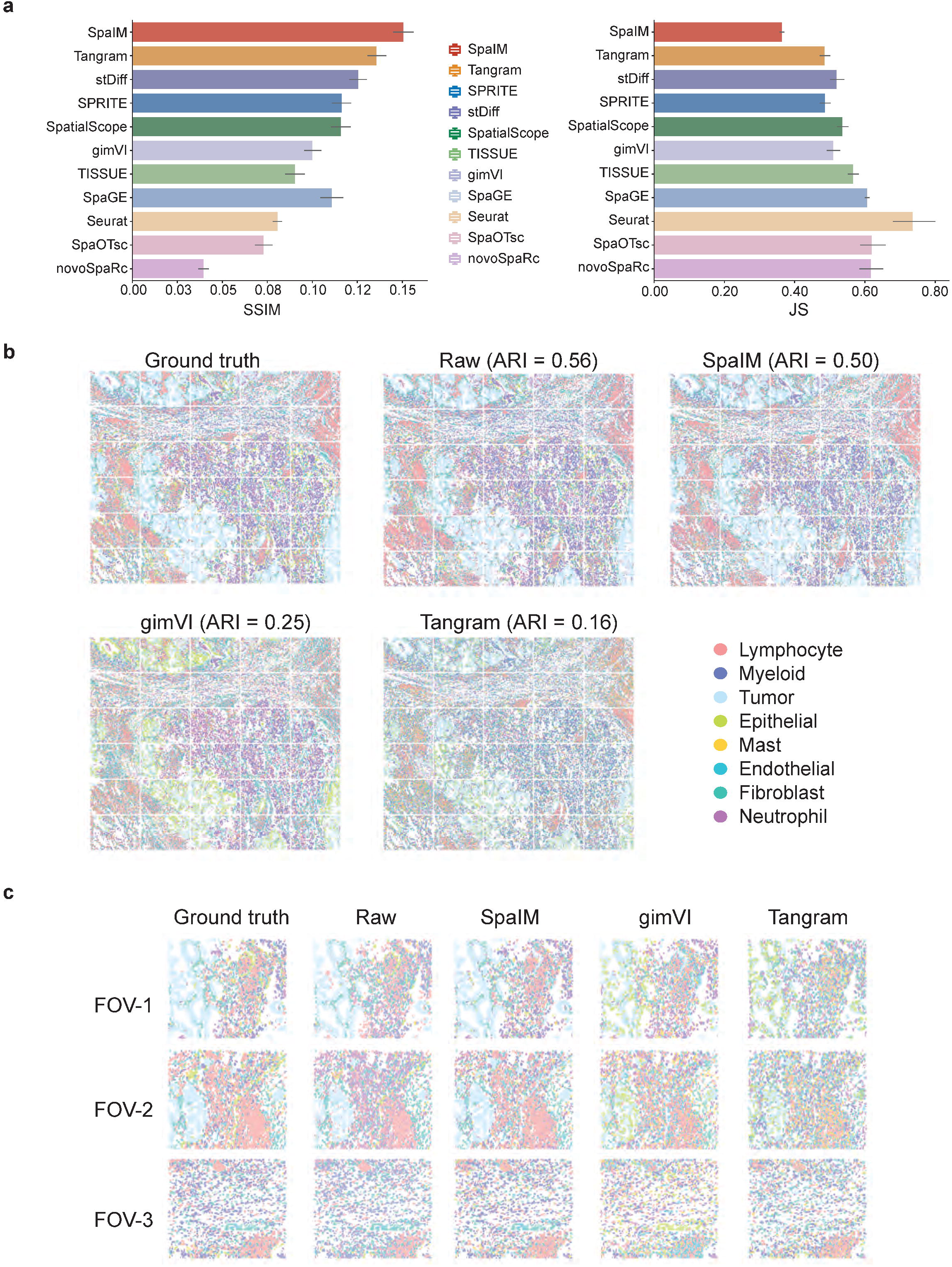
SpaIM facilities spatial domain detection. **a** Benchmarking results on the NanoString CosMx spatial transcriptomics dataset (Lung5-rep3), using evaluation metrics including structural similarity index measure (SSIM) and Jaccard similarity (JS). **b** Spatial visualization of cell types in the whole slide. **d** Spatial visualization of cell types specific Field Of Views (FOVs).

We next evaluate the accuracy of spatial domain detection based on imputed gene expression data generated by different methods. **Fig.4a** shows the Adjusted Rand Index (ARI) scores of all 20 FOVs in the Lung5-rep3 dataset. Notably, SpaIM achieves the highest accuracy in identifying spatial domains corresponding to different cell types (ARI = 0.50), closely approximating to the ground truth and raw data (ARI = 0.56). In contrast, Tangram and gimVI produced significantly lower ARI scores of 0.16 and 0.25, respectively. Further analysis at the individual FOV level (**Fig.4a**) demonstrates that SpaIM consistently aligns with the ground truth in distinguishing cellular identities. SpaIM accurately reveals a continuous tumour region infiltrated with dispersed immune cells. Conversely, gimVI and Tangram exhibit errors in cellular structure identification, often generating fragmented or blended cell type regions. For example, in FOV-1, both gimVI and Tangram misclassify tumour cells as epithelial cells. In FOV-2, lymphocytes are incorrectly identified as neutrophils, and in FOV-3, gimVI fails to differentiate lymphocytes, misclassifying them as fibroblasts. These errors result in poor delineation of spatial heterogeneity. These findings highlight SpaIM’s superior performance in leveraging imputed gene expression to achieve precise spatial domain identification compared to alternative methods.

SpaIM demonstrates remarkable ability in imputing unmeasured spatial gene expression data, providing biologically meaningful insights. As illustrated in the UMAP plots (**Fig.5a**), SpaIM effectively imputes cell type-specific gene markers, maintaining consistent expression patterns within their respective cell types (**Fig.5a**). For instance, the endothelial marker gene *MME*^43^ exhibits uniform high expression in endothelial cells and low expression in other cell types when imputed by SpaIM. In contrast, Tangram’s imputed data shows sporadic MME expression in endothelial cells and pervasive expression in non-endothelial cell types. Other SpaIM-imputed cell type-specific markers, *CD1C*^44, 45^ and *DAG1*^46^ are enriched in lymphocyte- and myeloid-dense regions, respectively, while the tumor-specific gene *MYCN*^47^ is strongly and uniformly expressed in tumor regions. These distinct patterns are not observed in Tangram’s imputed data (**Fig.5a**). These results highlight SpaIM’s reliability in imputing unmeasured gene expression, enabling more accurate spatial cellular characterization.

**Fig. 5.**
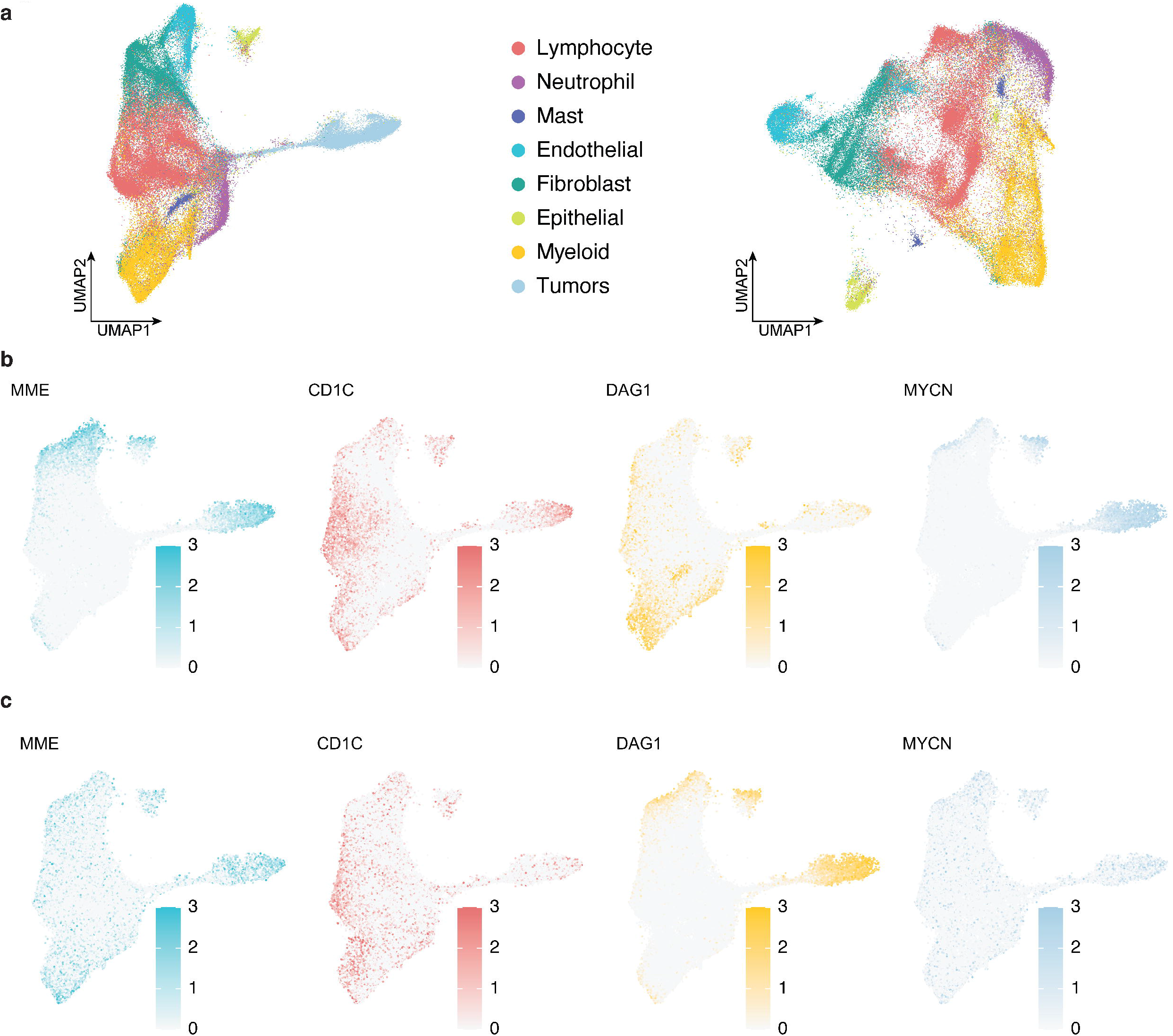
SpaIM reliably recovers unmeasured genes. **a** UMAP visualizations of all cell populations and nontumor populations. **b** UMAP visualization of the imputation of unmeasured marker genes by SpaIM. **c** UMAP visualization of the imputation of unmeasured marker genes by Tangram.

### SpaIM accurately imputes spatial gene expression across diverse ST platforms

To further evaluate SpaIM’s performance, we conduct comprehensive experiments across multiple spatial transcriptomics (ST) datasets, profiling both sequencing-based and imaging-based platforms. Specifically, we collect 21 Visium datasets encompassing a range of tissues, including breast, prostate, kidney, and brain, from humans, mice, and zebrafish (**Supplementary Table 1**). Considering the diverse data quality and characteristics across datasets, we employ ranked performance for a fair comparison across different methods^48^. The benchmarking results of different methods on 10x Visium datasets are shown in **Fig.6a** and **Supplementary Fig.2a. Fig.6a** highlights that SpaIM consistently achieved the highest performance across these datasets, outperforming the second-best model by more than 13% in ACC.

**Fig. 6.**
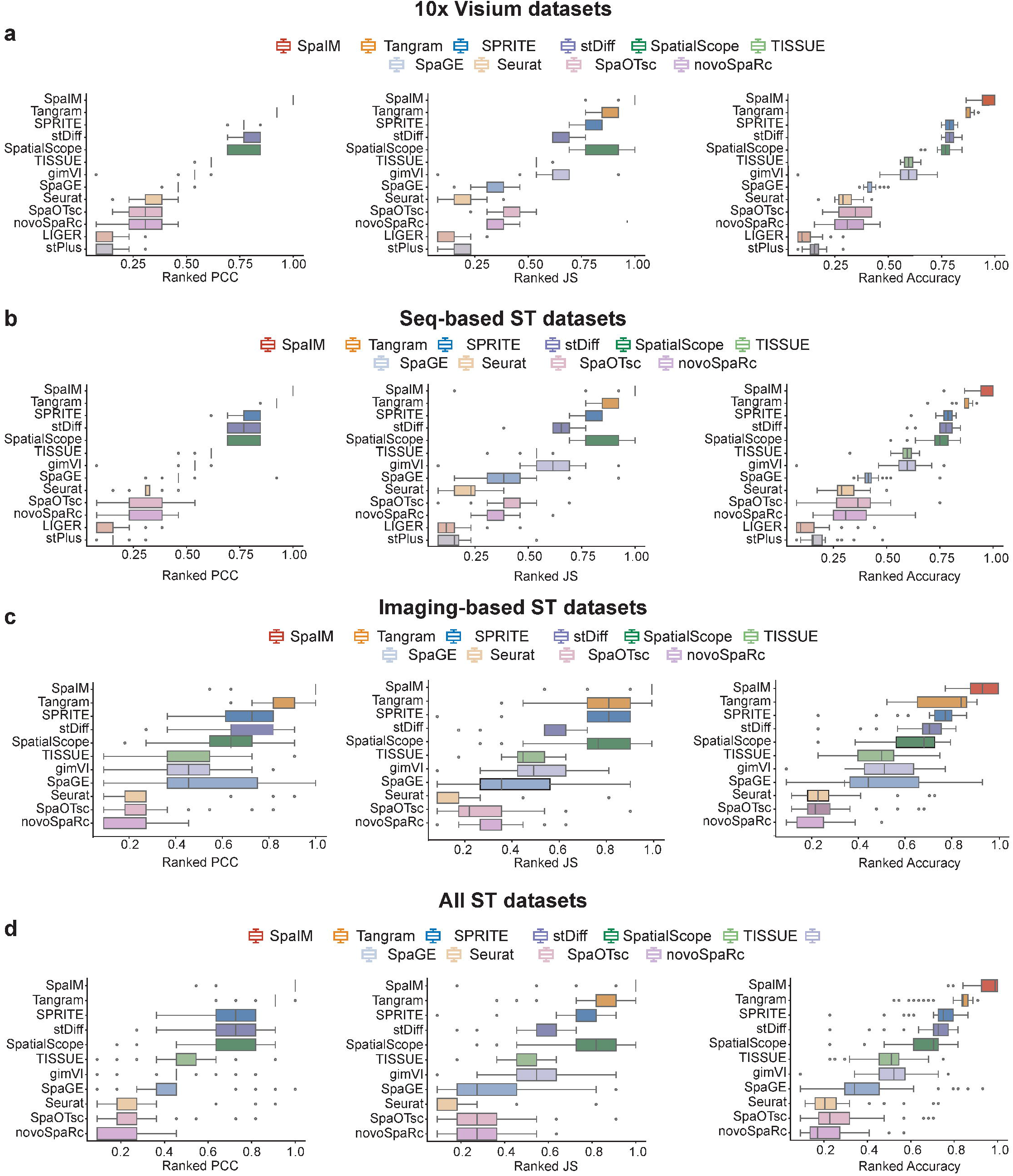
Gene imputation performance across diverse spatial transcriptomics (ST) datasets. Comparison results across (**a**) 21Visium ST datasets; (**b**) 28 sequencing-base datasets; (**c**) 25 imaging-based ST datasets; (**d**) total 53 datasets profiled by diverse ST technologies, using ranked Pearson correlation coefficient (PCC), Jaccard similarity (JS), and Accuracy score.

Moreover, we expand the evaluations to include other sequencing-based platforms such as Slide-seq, Slide-seq V2, Seq-scope, and HDST. This expanded sequencing-based ST data collection includes totally 28 datasets, with their paired scRNA-seq data from the same tissue type (**Supplementary Table 1**). The benchmarking results on these datasets are illustrated in **Fig.6a** and **Supplementary Fig.2a**. Notably, SpaIM continues to exhibit superior performance across all sequencing-based datasets than other methods. Specifically, SpaIM has higher PCC (1.0±0.0), JS (0.96±0.16), and ACC (ACC: 0.97±0.06) than other methods such as Tangram (PCC: 0.92±0.02, JS: 0.87±0.05, ACC: 0.87±0.04), gimVI (PCC: 0.51±0.13, JS: 0.60±0.17, ACC: 0.57±0.15), and stDiff (PCC: 0.77±0.06, JS: 0.65±0.07, ACC: 0.77±0.05).

We also showcase the exceptional gene imputation performance of SpaIM across 25 imaging-based ST datasets, including platforms such as seqFISH, seqFISH+, MERFISH, STARmap, ISS, and osmFISH (**Supplementary Table 1**). These datasets typically exhibit limited gene coverage and signals. Based on these datasets, we conduct experiments using SpaIM and other methods, with results showing in **Fig.6a** and **Supplementary Fig.2a**. SpaIM achieves more accurate predictions with higher PCC (0.97±0.11), JS (0.95±0.12), and ACC (ACC: 0.92±0.08) than other methods such as Tangram (PCC: 0.86±0.10, JS: 0.77±0.19, ACC: 0.78±0.13), gimVI (PCC: 0.47±0.18, JS: 0.48±0.18, ACC: 0.50±0.17), and stDiff (PCC: 0.67±0.16, JS: 0.59±0.13, ACC: 0.68±0.14).

A comprehensive evaluation across 53 datasets, spanning both sequencing- and imaging-based platforms, highlights SpaIM’s exceptional performance. These datasets, encompassing diverse data quality and characteristics, provide a robust benchmark for comparison. As illustrated in **Fig.6a** and **Supplementary Fig.2a**, SpaIM consistently outperforms other methods, achieving an average ACC of 0.95 ± 0.07. In contrast, Tangram (ACC: 0.81 ± 0.10), gimVI (ACC: 0.50 ± 0.15), and stDiff (ACC: 0.71 ± 0.10) demonstrate comparatively lower performance. This extensive assessment underscores SpaIM’s reliability and robustness, establishing it as a state-of-the-art tool for spatial transcriptomics across diverse datasets.

## DISCUSSION

In this paper, we introduce SpaIM, a novel style transfer learning model designed to impute unmeasured spatial gene expressions. SpaIM adopts a novel strategy that reconceptualizes spatial gene expressions into dataset-agnostic gene contents and dataset-specific styles. The SpaIM model comprises the Recursive Style Transfer (ReST)-based ST autoencoder and the ReST-based ST generator. Central to SpaIM is its dual-task style-transfer learning approach: the ST autoencoder disentangles ST gene expression patterns into content and style using scRNA-seq as a reference, while the ST generator transfers learned style to infer gene expressions in ST data from scRNA-seq inputs. Both components share a common decoder and are jointly trained with a loss function based on overlapping genes between ST and scRNA-seq data, enhancing the model’s ability to capture and interpolate gene expression relationships. SpaIM effectively integrates spatial data styles with the content from scRNA-seq data to accurately recover masked gene expressions, demonstrating its capability in imputing unmeasured genes in ST data, particularly in imaging-based ST that typically lack comprehensive measurements.

Compared to existing methods, SpaIM offers superior gene imputation performance and distinguishes itself from others. We reason that other methods generally try to locally align scRNA-seq with ST for spatial gene imputation, while overlooking the stylistic differences between the two types of data. SpaIM innovatively redefines gene imputation by decoupling of both scRNA-seq and ST datasets into dataset-agnostic contents and dataset-specific styles.

This strategic decoupling allows SpaIM to better recognize and utilize both the commonalities and unique characteristics between scRNA-seq and ST data. Such clarity in data handling not only leads to more precise gene imputation but also improves the model’s ability to adapt to various ST data characteristics, thereby boosting SpaIM’s generalization capabilities and applicability across different from various platforms (i.e. sequencing-based and imaging-based). Such clarity of the SpaIM design provides better interpretability in its accurate predictions of incomplete gene expression data, positioning SpaIM as a leading method in the field. More importantly, SpaIM greatly enhances downstream analyses in spatial transcriptomics data, opening avenues for biological discovery. By accurately imputing missing gene expressions, SpaIM enables the identification of key ligand-receptor pairs and enhances differential expression analyses, allowing for precise identification of spatial cell types and the discovery of genes that vary across them. This powerful tool expands the scope of spatial data analysis and enhance biological insight derived from spatial transcriptomics.

In addition to its superior performance and technical advantages, SpaIM has potential areas for future improvement. First, SpaIM currently employs straightforward Multi-Layer Perceptron (MLP) layers, which may potentially benefit from more sophisticated designs such as graph transformer^49^ and the cutting-edge Mamba^50^. Second, as SpaIM utilizes independent initial input values for the ST generator, there is potential for improvement by developing customized styles specifically tailored to distinct datasets. Then the ST generator can better understand and adapt to the unique characteristics and nuances of different spatial datasets, thereby enhancing the accuracy of spatial gene imputation. Third, we anticipate improving the interpretability of the SpaIM model to provide deeper insights into missing gene expressions and explore the underlying mechanisms in tissue ecosystems. In summary, SpaIM represents a significant advancement in spatial gene expression imputation and is anticipated to facilitate biological discoveries and insights into complex tissues and diseases.

## MATERIALS AND METHODS

### Data Preprocessing

We include 28 datasets profiled from sequencing-based ST technologies including 10x Visium, and 25 datasets from imaging-based ST technologies, such as seqFISH+, MERFISH, and NanoString CosMx SMI. For each ST dataset of *G*_1_ genes and cells *C*_1_, we select a corresponding scRNA-seq dataset of *G*_1_ genes and *C*_2_ cells from the same tissue type as the reference, with *G*_1_ ≫ *G*_2_. Both raw count data are preprocessed with log1p-transformed. The processed ST and SC gene expression data is denoted as 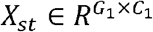 and 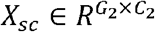,respectively. The individual cell-level information in*X*_*sc*_ carries significant noise due to dropouts, duplets, inaccurate cell segmentation, and RNA degradation. Instead of representing the gene expression of a gene with all *C*_2_ cells, we employ Leiden clustering to group cells into *K* clusters and use the median expression of the gene in each cluster to represent the gene. Thus, we derive 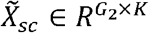 from 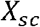.

The common genes shared by both ST and SC data are randomly split at the 80:20 ratio into the training set with *G* genes to train the model, and the validation set with *G* ^′^ genes to evaluate the performance. The trained model will then be used to infer the spatial gene expression of the genes that are only available in the SC dataset but not in the ST.

### The SpaIM model

SpaIM is a recursive style-transfer deep learning model with an ST autoencoder and an ST generator. The goal of SpaIM is to infer the expressions of genes that are only available in SC but not in the ST data, while keep the gene expression patterns as if they were measured using the ST platform.

#### 1) Recursive Style Transfer (ReST) layer

The fundamental component of both the ST autoencoder and the ST generator is the ReST layer with layer-wise fusion. As shown in **Supplementary Fig.3a**, the ‘th ReST layer is composed of a content encoder *C*^(*l*)^, a style *S*^(*l*)^ encoder, and a decoder *D*^(*l*)^ The ReST layer has two functionalities: encoding the received representations of content (*h*^(*l*)^) and style (*g*^(*l*)^)to latent representation *h*^(*l*+1)^ and *g*^(*l*+1)^, respectively, and output them, fusing the extracted latent content and style to the reconstructed data (*p*)^(*l*)^ and output it. The architecture of the ReST layer allows building recursive style transfer models through layer-wise feature extraction and fusion (**Fig.1a**). The ReST layer is a general style transfer with versatile uses. For example, if the same data is used as the input of content as well as the style, with appropriate loss functions, the ReST layer becomes an autoencoder to disentangle content and style. If a virtual style is used, with appropriate loss functions, the ReST layer becomes a generative model to impose a style to the content. The ST autoencoder and the ST generator use such design strategies.

#### 2) Spatial Transcriptomics (ST) Autoencoder

The ST autoencoder (**Fig.1a**) comprises multilayer ReST encoders. At each layer, the content encoder 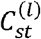 and the style encoder 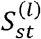 extract representation of the content and the style and a decoder *D*^(*l*)^ for layer-wise fusions of the learned content and style, with *l* = 0,1…,*L* representing different layers. Each encoder layer comprises a linear sub-layer, followed by a normalization sub-layer, and finally a ReLU sub-layer. The latent representation of the content in the ‘th layer is:

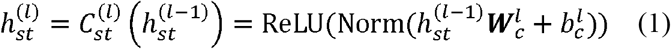

and the style representation is:

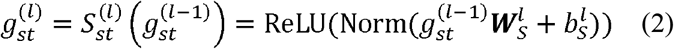

where 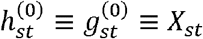, with 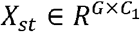 denoting the input spatial transcriptomics, *G* is the number of genes, and *C*_1_ is the number of cells.

The decoder layers fuse the latent representations of content and style layer by layer, starting from the layer *L*’th layer (e.g., the last layer) till the first layer (i.e., *l* =). The *k*-th decoded layer is:

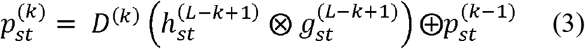

where ⊗ denotes the element-wise multiplication of the content and the style latent representation, ⊕ is the element-wise addition operator, *k*=1,2,…*L*, and 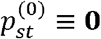.For example, when *L*=2

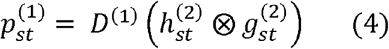

and

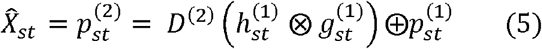

where 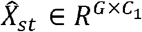 is the reconstructed ST data. The recursive architecture allows the latent representations of contents and styles as well as the reconstruction of the data at different resolutions, and the layer-wise fusion of the reconstructed data.

#### 3) Spatial Transcriptomics (ST) Generator

A similar architecture (**Fig.1a**) is used to generate ST data from SC data. Briefly, the latent representation of the content in the *l*’th layer is:

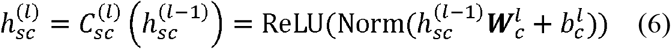

and the style representation is:

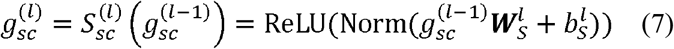

The genes in the input data varies in the training, validation, and inferring. During training, 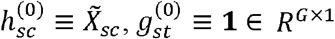.

The ST generator shares the decoder with the ST autoencoder. That is, the ST generator does not have its own decoder. Therefore, the *k*’th decoded layer is:

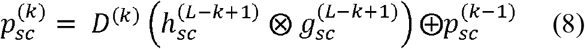

where 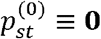.For example, when *L*=2.

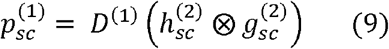

and

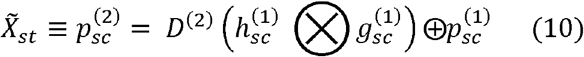

where 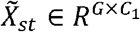 is the inferred ST data through style transfer and the final model output.

During testing, 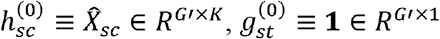, and 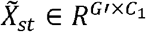 During inferring, the SC gene expression of any SC gene can be used as the input to generate the corresponding spatial gene expression for each cell in the ST data.

#### 4) Loss function

The loss function is composed of four components: i) the content loss, ii) the style loss, and iii) two reconstruction losses for the ST autoencoder and the ST generator, respectively. The content loss and the style loss are used to disentangle content and style. The style loss also enables style transfer. The ST autoencoder reconstruction loss is for learning an enhanced ST data, which is used by the ST generator reconstruction loss for accurately inferring spatial gene expressions from SC data.

### Content loss

Content loss is primarily utilized to optimize the similarity between the learned content at each encoding layer, aiding the model in preserving the essential information of the gene data. By minimizing the differences between content features, we ensure that the spatial gene data generated by the ST generator remains functionally consistent with the ST data. For the *L* layers in the model, the content loss (ℒ_*content*_) of the latent space was the summation of the mean square error (MSE) between the *l*-th content features from the spatial 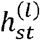 and scRNA dataset 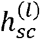:

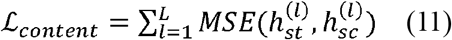

### Style Loss

Gram matrixes are used to represent the learned style at each layer, which is similar to the style representation in neural image style transfer frameworks^32^. The gram matrix for the spatial style extracted from the ST data and the SC data at the *l*-th layer are:

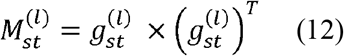

and

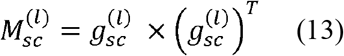

and the style loss is:

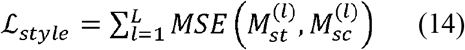

### Reconstruction Loss

The ST data reconstruction by the ST autoencoder is optimized by minimizing the difference between the original and the reconstructed ST data using cosine similarity:

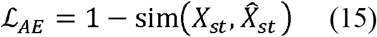

where the cosine similarity is defined as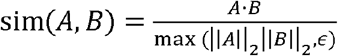, is a small positive number for robustness. Similarly, the ST generator reconstruction loss is defined as:

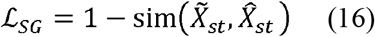

The overall loss

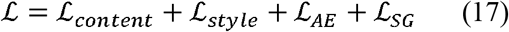

thus optimizes the model’s training process from the aspects of feature similarity, style transformation, and data accuracy. The number of layers and latent dimensions of SpaIM model are determined by grid-based fine-tuning across all datasets (**Supplementary Fig.3a-c**).

### Evaluation metrics

To accurately and comprehensively evaluate the performance of different models, we have employed several evaluation metrics in the model evaluation process, including the Pearson Correlation Coefficient (PCC), Structural Similarity Index (SSIM), Root Mean Square Error (RMSE), Jaccard Similarity (JS), and accurate score. These metrics assess the similarity and accuracy between the generated spatial gene data and the original data from different perspectives.

Pearson Correlation Coefficient (PCC): PCC measures the linear correlation between the generated spatial gene data and the original data. A higher PCC value indicates a stronger linear correlation between the generated and true data, indicating that the generated data maintains the basic trends and patterns of the original data well.

Structural Similarity Index Measure (SSIM): SSIM evaluates the structural similarity between the generated spatial gene data and the original data. It considers not only differences in brightness but also structural aspects of the data. Therefore, SSIM provides a more comprehensive assessment of the similarity between the generated and true data.

Root Mean Square Error (RMSE): RMSE measures the average error between the generated spatial gene data and the original data. A lower RMSE value indicates smaller differences between the generated and true data, implying higher prediction accuracy of the generated data.

Jaccard Similarity (JS): JS assesses the similarity, particularly focusing on the similarity of gene expression patterns, between the generated spatial gene data and the original data. It quantifies the overlap between two sets of genes, thus evaluating the consistency between the generated and true data.

Accurate score (ACC): ACC is a comprehensive metric that evaluates the overall accuracy of the generated spatial gene data compared to the original data. It provides an integrated measure of the aforementioned performance and utilizes the ranked relative values to assess the quality of the generated data in gene expression analysis tasks.

Ranked metrics: Using the same relative value ranking method as for ACC, we apply it to three other metrics, resulting in ranking Jaccard similarity (Ranked JS), ranking Pearson correlation coefficient (Ranked PCC), ranking structural similarity index measure (Ranked SSIM), and ranking root mean square error (Ranked RMSE). For all these metrics, higher values indicate better model performance.

### Implementation detail

During training, the model utilizes a learning rate of 0.001, runs for a maximum of 300 epochs. The experiments are performed on an Ubuntu 20.04 system equipped with 128GB of RAM and an NVIDIA GeForce RTX 3090 Ti GPU featuring 24GB of memory.

## FIGURE LEGENDS

**Supplementary Fig.1aBenchmarking evaluation between SpaIM and other methods. a** Comparisons between SpaIM and existing methods on the breast cancer dataset using Jaccard similarity (JS), root mean square error (RMSE), and Accuracy score. **b** Comparison results between SpaIM and existing methods on Lung9-rep1 dataset using Pearson correlation coefficient (PCC), root mean square error (RMSE), and Accuracy score. **c** Comparisons between SpaIM and existing methods on Lung5-rep3 using Pearson correlation coefficient (PCC), root mean square error (RMSE), and Accuracy score.

**Supplementary Fig.2aBenchmarking evaluation of imputation performance across diverse spatial transcriptomics (ST) datasets**. Comparison results across (**a**) 21Visium ST datasets; (**b**) 28 sequencing-base datasets; (**c**) 25 imaging-based ST datasets; (**d**) total 53 datasets profiled by diverse ST technologies, using evaluation metrics, i.e. ranked root mean square error (RMSE) and ranked structural similarity index measure (SSIM).

**Supplementary Fig.3a** (**a**) Illustration of the Recursive Style Transfer (ReST) layer. Grid-based hyper-parameter fine turning of latent dimensions (**b**) and the number of layers (**c**). In the boxplot, the center line, box limits and whiskers denote the median, upper and lower quartiles, and 1.5× interquartile range, respectively.

**Supplementary Table 1** Details of spatial transcriptomics data and single-cell RNA-sequencing data.

## CODE AVAILABILITY

All source codes and trained models in our experiments have been deposited at https://github.com/QSong-github/SpaIM.

## DATA AVAILABILITY

All datasets used in this study are publicly available. Data sources and details are provided in **Supplementary Table**

## ETHICS DECLARATIONS

## MATERIALS & CORRESPONDENCE

Correspondence and requests for materials should be addressed to BY, JS, or QS.

